# Transpirational water loss from flowers is low but unregulated

**DOI:** 10.1101/2024.06.28.600864

**Authors:** Adam B. Roddy, Jeroen D. M. Schreel, Dario Caminha Paiva, Ni Qin, Guo-Feng Jiang, Craig R. Brodersen, Kevin A. Simonin

**Affiliations:** Institute of Environment, Department of Biological Sciences, Florida International University, Miami, FL, USA; Plant Sciences Unit, Flanders Research Institute for Agriculture, Fisheries and Food (ILVO), B-9090 Melle, Belgium; Guangxi Key Laboratory of Forest Ecology and Conservation, Guangxi Colleges and Universities Key Laboratory for Cultivation and Utilization of Subtropical Forest Plantation, and State Key Laboratory for Conservation and Utilization of Subtropical Agro-Bioresources, College of Forestry, Guangxi University, Daxuedonglu 100, Nanning, Guangxi 530004, PR China; School of the Environment, Yale University, New Haven, CT, USA; Department of Biology, San Francisco State University, San Francisco, CA, USA

## Abstract

Flowers play a critical role in reproduction for most of the flowering plants, and maintaining flowers throughout their lifespan can require substantial resources, such as carbon and water. Increases in temperature and aridity due to climate change are shifting the atmospheric conditions experienced by flowers, potentially altering the costs of floral maintenance. However, little is known about floral physiology and the capacity of flowers to regulate water loss. Because many flowers have few, if any, stomata, flowers may not be able to curtail water loss when the atmospheric demand for water vapor is high. Here, we tested whether the surface conductance of of flower petals, tepals, and showy bracts responds dynamically to changes in the vapor pressure gradient driving water loss. We measure the responses of flower surface conductance (gs) to step changes in the vapor pressure gradient on nine species. Across species, gs was low among all species, and there was little, if any, response in gs to step changes in humidity. The lack of response in gs resulted in linear responses of transpirational water loss to variation in vapor pressure deficit. These results suggest that unusually hot, dry conditions could elevate water loss from flowers, leading to premature wilting and senescence, thereby shortening floral longevity.

## Introduction

Individual plants are capable of moving up to hundreds of kilograms of water from the soil to the atmosphere each day (Wullschleger *et al*. 1998), which influences local and global climate cycles (Boyce and Lee 2010; Boyce *et al*. 2010). As water exits the plant, almost all of it passes through the stomata on the leaf surface, which also regulate the exchange of CO_2_ between the leaf and the atmosphere (Hetherington and Woodward 2003). Because opening stomata to increase leaf surface conductance to CO_2_ exposes the wet leaf interior to the dry atmosphere and greater evaporative water loss, increases in net CO_2_ uptake from the atmosphere have required increases in the capacity to transport liquid water throughout the entire plant hydraulic pathway. In leaves, these increases have occurred predominantly through reducing the sizes of stomata on the leaf surface and increasing the packing densities of stomata and xylem veins (Brodribb and Feild 2010; Feild *et al*. 2011; Sack and Buckley 2016; Simonin and Roddy 2018). Because excessive water loss can cause declines in water content that impair physiological function, stomatal physiology–how rapidly stomata open and close in response to changes in atmospheric humidity and the rate of leaf water loss–is critical in regulating plant water balance (Mott and Parkhurst 1991; Brodribb and McAdam 2011; Lawson and Vialet-Chabrand 2019). Leaves, therefore, have evolved to coordinate the delivery of high rates of CO_2_ and water when light is available while also rapidly curtailing water loss when environmental conditions could lead to physiological impairment.

However, during certain time periods, leaves are not the only organs through which water escapes to the atmosphere. As plants transition into their reproductive phase, they allocate increasing amounts of resources to building and maintaining flowers and fruits (Bazzaz *et al*. 1987). Flowers function primarily to promote outcrossing, using visual, chemical, and thermal signaling to attract pollinators (Kevan 1975; Whitney *et al*. 2011; von Arx *et al*. 2012; van der Kooi *et al*. 2019; Dahake *et al*. 2022). Though they develop from the same shoot apical meristem as leaves and are often composed of laminar organs, flowers have experienced different selection from leaves due to their different functions. For example, the showy organs of the floral perianth are often heterotrophic and so do not need to maintain high fluxes of CO_2_ diffusion across their epidermises (Olson and Pittermann 2019; Roddy *et al*. 2019; Roddy 2019; but see Werk and Ehleringer 1983; Galen *et al*. 1993). As a result, these showy floral structures often have fewer stomata than their conspecific leaves (Lipayeva 1989; Blanke and Lovatt 1993; Roddy *et al*. 2016; Zhang, Carins Murphy, *et al*. 2018). In some cases, flowers may entirely lack stomata (Roddy *et al*. 2019), and some evidence suggests that even when stomata are present on flowers, they may not be able to close (Hew *et al*. 1980; Teixido and Valladares 2014). While having fewer stomata may limit maximum rates of water loss from flowers, the lack of stomata or the presence of non-functional stomata comes with the potential cost that water loss from flowers and the associated latent heat exchange cannot be regulated, potentially resulting in excessive transpiration during hot, dry atmospheric conditions (Blanke and Lovatt 1993; Teixido and Valladares 2014; Roddy 2019; Bourbia *et al*. 2020; Carins-Murphy *et al*. 2023). However, the temporal dynamics of water loss regulation from flowers has been poorly studied (Gleason 2018), despite the potentially large effects of the physiological costs of producing and maintaining flowers on flower function and evolution (Caruso *et al*. 2018).

Here, we tested whether the surface conductance (*g*_s_) of flower petals, tepals, and showy bracts responds dynamically to changes in the vapor pressure gradient driving transpirational water loss. In flowers lacking stomata, this surface conductance would be driven entirely by the cuticular conductance, while in flowers with non-functional stomata this surface conductance would be a combination of the stomatal conductance and the cuticular conductance. Even if floral stomata are functional, there may be so few stomata that cuticular conductance dominates the transpirational flux such that stomatal closure may have little effect on total transpiration rate. We hypothesized that because flowers have few stomata, transpiration rates would be relatively low compared to leaves but that flowers would be unable to rapidly regulate their *g*_s_ in response to changes in humidity. Therefore, transpirational water loss (*E*) from flowers was predicted to increase linearly with the vapor pressure difference (VPD) between the flower and the atmosphere.

## Results

Across species, floral *g*_s_ and *E* were relatively low compared to leaves, with measured g_s_ never exceeding 0.05 mol m^-2^ s^-1^ for any species. With the exception of two subtropical species, *P. platantha* and *B. blakeana*, g_s_ was typically below 0.02 mol m^-2^ s^-1^ for all species.

Across species, we found little, if any, responses of *g*_s_ to step changes in VPD over 20 minutes of measurement, despite over 1 kPa changes in VPD (Figure 1). If anything, there was a slight increase in *g*_s_ at higher VPD, though this effect was small (Figure 2). In some species, such as *M. erythrochlamys* and *T. erecta, g*_s_ did decline slowly after the step change to high VPD and increased slightly after the return back to low VPD. However, given the low fluxes, these changes in surface conductance were miniml. For example, in *T. erecta*, the decline of *g*_s_ over almost 20 minutes at a VPD of 2.5 kPa was less than 0.01 mol m^-2^ s^-1^.

**Figure 1.**
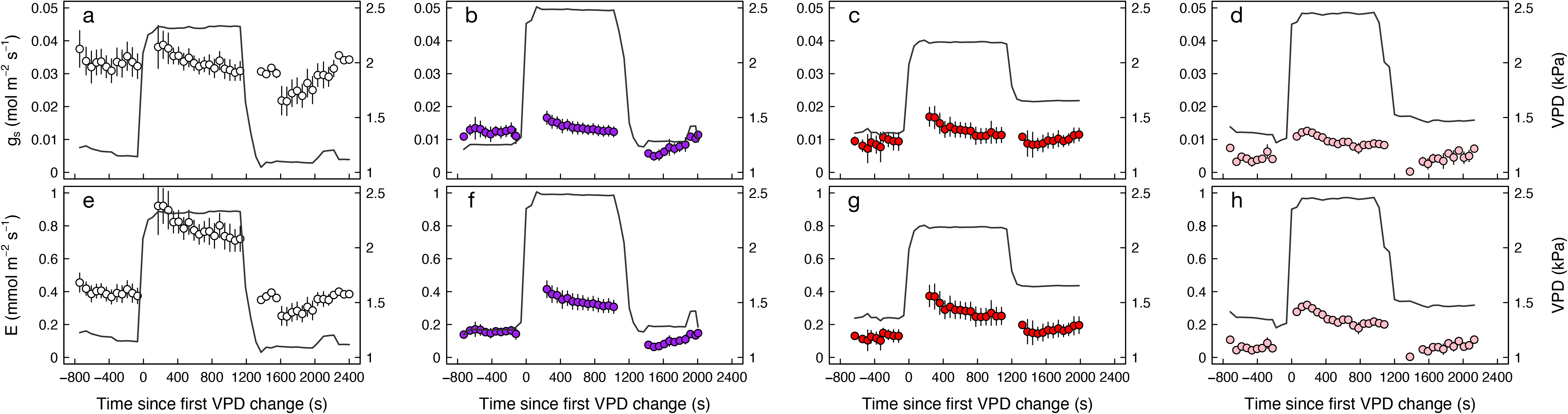
Time series of *g*_s_ (top row) and E (bottom row) responses to step changes in the flower-to-air vapor pressure deficit (solid line) driving transpiration. (a,e) *Portlandia platantha*, (b,f) *Thunbergia erecta*, (c,g) *Megaskepasma erythrochlamys*, (d,h) *Bougainvillea spectabilis*. Points and error bars represent means ± s.e. of 60-second bins across replicate flowers. Inset images show stomata for each species.

**Figure 2.**
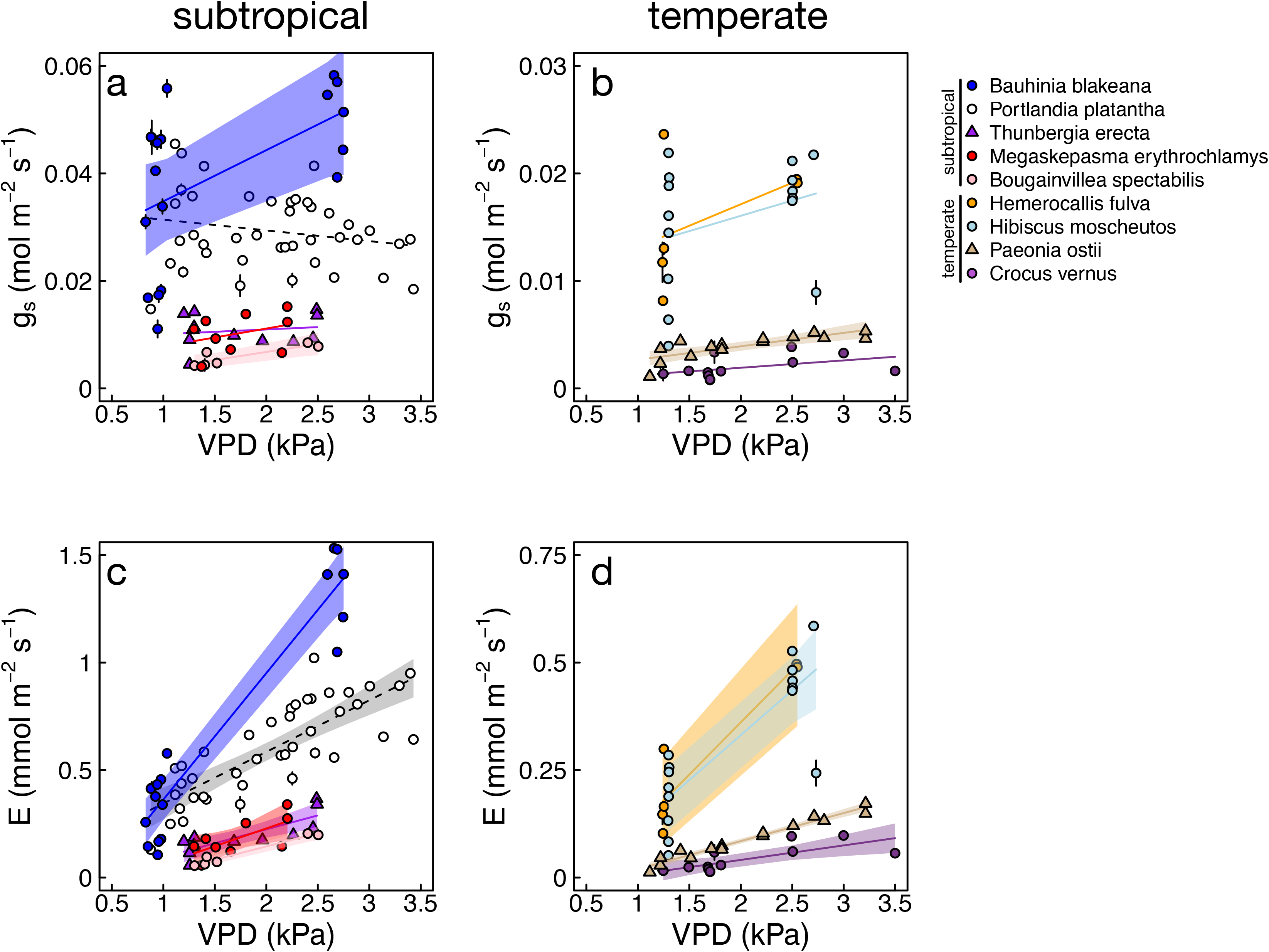
Relationships between vapor pressure deficit (VPD) and (a, b) *g*_s_ and (c, d) *E* for flowers of each species. Species are split according to habitat: (a, c) subtropical species, (b, d) temperate species. Note that the scales on the y-axes differ between plots for temperate and tropical species. Each point represents the final 60-second average of *g*_s_ or *E* for each sample at each VPD. Lines and shading represent linear regressions and 95% confidence intervals. Note that all linear regression are plotted, but 95% confidence intervals are plotted for species for which the regressions were statstically significant at α = 0.05. Points are colored according to species.

In contrast to *g*_s_, there were large, immediate effects of VPD changes on *E*. In all species, step increases in VPD resulted in immediate increases in *E*, which were not ameliorated by time due to the limited g_s_ responses. Similarly, step decreases back to low VPD resulted in large, immediate declines in *E* that remained largely stable due to the limited changes in *g*_s_ over time. For example, in *P. platantha*, which had the highest measured *g*_s_, a step change of over 1 kPa in VPD resulted in an increase of *E* from approximately 0.4 mmol m^-2^ s^-1^ to 0.8 mmol m^-2^ s^-1^ while *g*_s_ remained between 0.03 and 0.04 mol m^-2^ s^-1^ (Figure 1).

Pooling the final, stable measurements of *g*_s_ and *E* at each VPD for each species showed the overall effects of VPD on *g*_s_ and *E*. For most species, there was no relationship between VPD and *g*_s_ across a more than 1 kPa range in VPD (Figure 2, Table 1). However, *P. ostii, B. spectabilis*, and *B. blakeana* exhibited significant, positive relationships between VPD and *g*_s_, suggesting that measured *g*_s_ is higher at higher VPD. Although not significant, the relationships between *g*_s_ and VPD for all other species except for *P. platantha* were also positive (Table 1). The lack of responsiveness of *g*_s_ to VPD resulted in significant, positive relationships between VPD and *E* for all species, although the magnitude of this effect (i.e. the slope) varied among species (Figure 2; Table 1).

**Table 1.** List of sampled species, location sampled, their total stomatal density (Ds, the sum of abaxial and adaxial densities in units of mm^-2^), and their coefficients and summary statistics of linear regressions between *g*_s_ and *E* as a function of flower-to-air vapor pressure difference (VPD). FBG = Fairchild Tropical Botanic Garden, Miami, USA; MBG = Marsh Botanic Gardens, New Haven, USA; AA = Arnold Arboretum, Jamaica Plain, USA; GXU = Guangxi University, Nanning, China. Asterisks indicate statistically significant relationships: ns = not significant, \**P* < 0.05, \*\**P* < 0.001.

| Taxon | Family | Location | $D_s$ | $g_s$ | | $E$ | |
| --- | --- | --- | --- | --- | --- | --- | --- |
| | | | | slope | $R^2$ | slope | $R^2$ |
| *Bauhinia blakeana* Dunn | Fabaceae | GXU | $12.8 \pm 1.5$ | 0.095 | 0.29* | 0.59 | 0.91** |
| *Portlandia platantha* Hook.f. | Rubiaceae | FBG | $4.2 \pm 1.3$ | -0.002 | 0.04<sup>ns</sup> | 0.24 | 0.64** |
| *Thunbergia erecta* (Benth.) T.Anderson | Acanthaceae | FBG | $7.1 \pm 2.29$ | 0.001 | 0.03<sup>ns</sup> | 0.13 | 0.66** |
| *Megaskepasma erythrochlamys* Lindau | Acanthaceae | FBG | $30.1 \pm 7.6$ | 0.003 | 0.11<sup>ns</sup> | 0.17 | 0.49* |
| *Bougainvillea spectabilis* Willd. | Nyctaginaceae | FBG | $11.9 \pm 2.1$ | 0.003 | 0.75* | 0.12 | 0.96** |
| *Hemerocallis fulva* (L.) L. | Asphodelaceae | MBG | 0 | 0.004 | 0.21<sup>ns</sup> | 0.24 | 0.86** |
| *Hibiscus moscheutos* L. | Malvaceae | AA | NA | 0.003 | 0.11<sup>ns</sup> | 0.21 | 0.67** |
| *Paeonia ostii* T.Hong & J.X.Zhang | Paeoniaceae | AA | 0 | 0.001 | 0.62** | 0.07 | 0.96** |
| *Crocus vernus* (L.) Hill | Iridaceae | MBG | NA | 0.001 | 0.20<sup>ns</sup> | 0.03 | 0.55** |

## Discussion

While having fewer stomata or non-functional stomata on perianth surfaces would result in lower maximum transpiration, our results highlight that having few stomata limits the capacity of flowers to regulate their surface conductance. Variation in the total water flux from a flower, therefore, is driven entirely by the vapor pressure difference driving evaporation across the epidermis and cuticle. At constant VPD, interspecific differences in floral transpiration rates are due, therefore, to differences in the residual conductance (*g*_res_, or, more commonly, the minimum conductance, *g*_min_) across the epidermis and through non-functional stomata. Under unusually hot, dry atmospheric conditions, the failure to limit water loss can cause flowers to wilt, shortening their lifespan and hindering their capacity to attract pollinators (Teixido and Valladares 2014; Teixido and Valladares 2015; Roddy *et al*. 2018; Bourbia *et al*. 2020; Carins-Murphy *et al*. 2023). When there are few stomata, selection to further reduce *g*_s_ of flowers would likely be focused on the cuticle, the waxy, hydrophobic covering on the epidermis. Although the petal *g*_s_ to water vapor was very low for most species studied here, broader surveys of minimum epidermal conductance on flowers has shown that it is variable among species and typically higher than that of conspecific leaves (Roddy *et al*. 2016, 2023). Though low, transpiration from flowers can be important to multiple aspects of flower functioning, including maintaining cool floral temperatures to prevent overheating (Patiño and Grace 2002; Roddy 2019) and helping to guide pollinators (von Arx *et al*. 2012; Sinha *et al*. 2022; Dahake *et al*. 2022). Therefore, functional demands beyond just transpiration rate may determine the optimal *g*_s_ for flowers.

These results illuminate possible changes in the physiology of floral stomata. The lack of a significant response of surface conductance to changes in the flower-to-air VPD are consistent with previous measurements on tropical orchids (Hew *et al*. 1980) and *Cistus* flowers growing in the hot Mediterranean summer (Teixido and Valladares 2014). These *Cistus* flowers also showed linearly increasing transpiration with increasing flower-to-air VPD. However, our results and those of Teixido *et al*. (2014) differ from results for *Magnolia* and *Calycanthus* flowers (Feild *et al*. 2009; Roddy *et al*. 2018). In these two magnoliid species, diurnal measurements of flower gas exchange showed that *g*_s_ varies throughout the day. Additionally, VPD responses measured on *Calycanthus occidentalis* revealed that floral *g*_s_ declines in response to increasing VPD, albeit not as rapidly or as effectively as leaf *g*_s_ (Roddy *et al*. 2018). The differences between our results and those for *Magnolia grandiflora* and *C. occidentalis* are not due solely to stomatal density, as stomata were present on *T. erecta, P. platantha, M. erythrochlamys, B. speciosa*, and *B. blakeana*. Nonetheless, flowers of these species exhibited limited capacity to regulate their surface conductance in response to large, rapid changes in VPD. Though completely lacking stomata undoubtedly indicates an inability to regulate flower surface conductance, many flowers that have stomata may still be unable to regulate their surface conductance, suggesting differences in stomatal physiology per se and not only anatomy. One potential mechanism is that flowers lack the ability to rapidly synthesize the hormone abscisic acid, which is responsible for rapid stomatal closure in angiosperm leaves (Brodribb and McAdam 2011; Zhang, Sussmilch, *et al*. 2018).

The response of surface conductance and transpiration rate to changes in VPD measured here have important implications for flower water balance. The relatively low transpiration rates, even under high VPD, allows for liquid water delivery rates–and the associated vein densities in flowers–to be low (Feild *et al*. 2009; Roddy *et al*. 2013; Roddy *et al*. 2016; Zhang, Carins Murphy, *et al*. 2018). Even though most of the water entering flowers is delivered throughout anthesis and not during earlier bud development (McMann *et al*. 2022), the overall low transpiration rates measured here suggest that floral hydraulic physiology may not operate under steady-state conditions typically employed by C3 leaves. Flowers have high hydraulic capacitance that would minimize declines in turgor presure even as water content declines (Roddy *et al*. 2019; An *et al*. 2023). A large hydraulic capacitor–likely due to the lower modulus of elasticity of the petal’s mesophyll and epidermal cells–would allow for large changes in cell volume (due to water loss) without substantial changes in the water potentials that drive liquid water flow through the xylem. Thus, even though transpiration rate may vary dynamically with changes in atmospheric conditions and flower temperature, the flux rates of water into flowers may be relatively low and constant because the flower-stem water potential gradient may be buffered from rapid changes due to high hydraulic capacitance. These dynamics would represent non-steady state physiology that is common when water content and hydraulic capacitance are high, such as occurs in flowers (Simonin *et al*. 2013; Roddy *et al*. 2023). During the day, when leaf surface conductance is higher than flower surface conductance, flowers may exhibit significantly longer water residence times than their conspecific leaves (Roddy *et al*. 2018). However, under conditions of low transpiration (e.g. at night or during droughts), leaf stomatal closure can reduce leaf surface conductance to be lower than flower surface conductance, resulting in flowers having relatively shorter water residence times than leaves, which could lead flowers wilting earlier than leaves (Roddy *et al*. 2023; Carins-Murphy *et al*. 2023).

Although the floral transpiration rates measured here were relatively low, at the canopy scale, flower water loss may be a significant contribution to whole-plant resource use during periods of flowering. The relative amounts of water lost from leaves and flowers would depend on the environmental conditions during flowering, the differences in surface conductances of leaves and flowers, and the differences in total leaf and flower evaporative surface areas on the whole plant (Blanke and Lovatt 1993; Teixido and Valladares 2014; Roddy *et al*. 2018; McMann *et al*. 2022). At the whole-plant level, water loss from flowers can be comparable to leaves and high enough to lower leaf water potential and suppress leaf gas exchange (Galen *et al*. 1999; Lambrecht and Dawson 2007; Lambrecht *et al*. 2011; Roddy and Dawson 2012; Lambrecht 2013). Furthermore, flowers are often positioned at the edge of plant crowns and above leaves, which may lead to higher light interception by flowers than leaves. In this scenario, flowers may have elevated temperatures and increased respiration rates while also shading leaves and suppressing leaf gas exchange rates (Teixido and Valladares 2014). In extreme cases, this effect of leaf shading and the added respiratory load of maintaining flowers can cause entire ecosystems to switch from being net carbon sinks to being net carbon sources (Sonnentag *et al*. 2011). By extension, water loss from flowers may also comprise a substantive contribution to ecosystem water balance during flowering periods, though this has received little attention.

Increasing evidence is showing that climate change is causing many plant species to invest more resources into flowering and reproduction (Wright and Calderon 2006; Anderegg *et al*. 2021; Bishop *et al*. 2023). Though reproduction has often been overlooked by ecophysiologists and biogeochemists as a relatively small component of biogeochemical cycles (Gleason 2018), higher investments in reproduction mean that plant reproductive structures may become a larger component of terrestrial biogeochemical cycling of carbon, water, and nutrients. That transpiration from flowers is unregulated and increases linearly with VPD reiterates the importance of flowers to whole-plant and ecosystem biogeochemical cycling. While the lack of physiological measurements on reproductive organs seriously hinders our knowledge of the role of reproduction in plant resource allocation, our results for water loss from flowers suggest that the water dynamics of flowering may be relatively easy to predict with some simple data. That surface conductance was unchanged with increasing VPD suggests that minimum epidermal conductance (*g*_min_) may be the primary physiological trait regulating water loss from flowers even under well-watered conditions. In leaves, *g*_min_ is thought to be critical to leaf drought tolerance because it determines how rapidly a leaf desiccates under water-limiting conditions that cause stomatal closure (Kerstiens 1996; Duursma *et al*. 2019). Because *g*_min_ can now be rapidly and cheaply measured (Billon *et al*. 2020), increasing the diversity of flower *<u>g</u>*_min_ sampling would provide powerful information about the water dynamics of individual flowers (Roddy *et al*. 2021; Roddy *et al*. 2023). Further, our results suggest that combining measurements of *g*_min_ with measurements of VPD would provide reasonable estimates of individual flower water use that could be scaled up to the whole canopy level with estimates of flower abundance.

## Methods

### Plant material

We sampled both temperate and subtropical plants growing at the Arnold Arboretum (Jamaica Plain, MA, USA), Marsh Botanical Gardens (New Haven, CT, USA), Fairchild Tropical Botanic Garden (Miami, FL, USA), and Gunagxi University campus (Nanning, Guangxi Province, China). We measured flowers from multiple individuals of each of nine species, except for *Paeonia ostii*, for which there were only two individuals available (Table 1). Species were chosen based primarily on the spatial constraints of enclosing floral organs in a gas exchange cuvette without damaging the flower or enclosing other floral organs that may introduce artifacts. For example, small flowers could have been measured by enclosing the entire flower in the chamber, but if there were nectar or water deep within the flower, it would have introduced artificially high transpiration rates. Therefore, we focused on species with laminar organs that could reasonably fit inside the cuvette. For species measured at the Arnold Arboretum, we excised flowering shoots in the early morning, immediately recut them underwater at least 25 cm away from the first cut, and transported the shoots in water back to the laboratory for measurements. For species measured at Marsh Botanical Gardens and Fairchild Tropical Botanic Garden, flowers remained on the plant during gas exchange measurements, which were performed in the morning.

### Responses of surface conductance (g_s_) to VPD

Gas exchange responses to vapor pressure deficit were measured using an LI-6800 (LiCor Biosciences, Lincoln, NE, USA) portable gas exchange system. Individual petals or tepals were enclosed in the gas exchange cuvette. In a few cases, small flowers required that multiple petals or tepals needed to be enclosed in the cuvette in order to prevent damage to other adjacent petals. We were sure to exclude sepals, which are often green and photosynthetic. For all measurements, reference CO_2_ concentration was held constant at either 415 or 420 ppm, and internal PAR was kept constant at 1000 μmol m^-2^ s^-1^. Data were logged every 30-60 s.

For measurements made in the laboratory, the LI-6800 was set to control petal temperature at 25{ º}C, but for measurements made in the field the LI-6800 was set to control chamber air temperature slightly higher than ambient air temperature to prevent condensation. Thus, temperature varied among samples depending on the ambient atmospheric conditions. Because fluxes were so low from flowers, there was relatively little latent heat loss, meaning that petal temperature was very stable even though petal temperature was not being controlled directly by the gas exchange system. The gas exchange system was set to control either VPD or, to reduce competition between system control loops, air saturation deficit. Initially these were set to 1.1-1.5 kPa, depending on the sample, and held for at least 10 minutes during which time surface conductance was stable.

A step increase to approximately 2.5 kPa was held for at least 20 minutes, followed by a decrease back to 1.2-1.5 kPa, with the exact value varying among samples but constant within samples. For some samples, the initial VPD started high and was reduced, followed by an increase, in order to test whether there was an effect of increasing versus decreasing VPD. For some samples, we measured gas exchange at multiple random VPDs in order to sample across a broader set of VPDs. We observed no qualitative differences in the responses of *g*_s_ and *E* to any of these three protocols, and so they were pooled ion analyses. Immediately after gas exchange measurements were completed, we carefully marked the areas of the petals that were enclosed inside the cuvette, excised these regions, and scanned them with a flatbed scanner to measure the enclosed one-sided projected surface area, which was used to recalculate gas exchange fluxes.

### Data analysis

To analyze the time courses of *g*_s_ responses to VPD, we pooled samples of each species that were measured under similar protocols (i.e. similar durations at each VPD step) and aligned them temporally to the first VPD step change. We then calculated means and standard errors across samples within 60-s bins. For presentation, we show these means ± s.e. for *g*_s_ and *E*, but for clarity we omit the s.e. around VPD.

For characterizing the effects of VPD on *g*_s_ and *E*, we calculated the mean ± s.e. of the last three stable point measurements at a given set of chamber conditions for each sample. Linear regression was used to determine the effect for each species of VPD on both *g*_s_ and *E*. All analyses were conducted in the R Statistical Software (v. 4.1.2; (Wickham 2017; R Core Team 2018; Vaughan and Dancho 2022; Wickham and Bryan 2023).

## Acknowledgments

This work was supported by a Gaylord Donnelley Fellowship from the Yale Institute for Biospheric Studies and a Jewitt Prize from the Arnold Arboretum of Harvard University to A. B. R. and by grants from the Yale Institute of Biospheric Studies and the US National Science Foundation (CMMI-2029756) to A. B. R. and C. R. B.

## Author Contributions

A. B. R. conceptualized the project. A. B. R., J. D. M. S., D. C. P., C. R. B., and K. A. S. designed the sampling. A. B. R., J. D. M. S., D. C. P., N. Q., and G.-F. J. collected the data. A. B. R. analyzed the data and wrote the first draft of the manuscript. All authors edited the manuscript.

